# Reference-dependent preferences arise from structure learning

**DOI:** 10.1101/252692

**Authors:** Lindsay E. Hunter, Samuel J. Gershman

**Affiliations:** Department of Psychology and Princeton Neuroscience Institute, Princeton University; Department of Psychology and Center for Brain Science, Harvard University

## Abstract

Modern theories of decision making emphasize the reference-dependency of decision making under risk. In particular, people tend to be risk-averse for outcomes greater than their reference point, and risk-seeking for outcomes less than their reference point. A key question is where reference points come from. A common assumption is that reference points correspond to expectations about outcomes, but it is unclear whether people rely on a single global expectation, or multiple local expectations. If the latter, how do people determine which expectation to apply in a particular situation? We argue that people discover reference points using a form of Bayesian structure learning, which partitions outcomes into distinct contexts, each with its own reference point corresponding to the expected outcome in that context. Consistent with this theory, we show experimentally that dramatic change in the distribution of outcomes can induce the discovery of a new reference point, with systematic effects on risk preferences. By contrast, when changes are gradual, a single reference point is continuously updated.

## Introduction

When people make decisions under risk (e.g., accepting or rejecting a gamble), their proclivity for risk depends critically on whether the outcome of the gamble is perceived as a gain or a loss [1]. If the outcome is perceived as a gain, most people will require an additional incentive (the risk premium) relative to a risk-neutral decision maker in order to accept the gamble, indicating that people are risk-averse for perceived gains. In contrast, most people are risk-seeking for perceived losses. The notion of “perception” is important here, because an objective gain may be perceived as a loss if it is less than expected (e.g., when one receives a surprisingly small raise), and likewise an objective loss may be perceived as a gain if it is greater than expected (e.g., a surprisingly inexpensive ticket). The expectation thus acts as a reference point for subjective valuation.

Modern theories of decision making have sought to formalize the concept of an expectation-based reference point and how it changes based on experience. In an influential line of work, Kőszegi and Rabin [2, 3] proposed that reference points reflect rational expectations based on recent outcomes (see also [4, 5]). In support of this theory, contestants on the TV game show “Deal or No Deal” were more likely to make risky choices when, upon receiving information that suggested they would take home less money than they expected [6], recapitulating results from laboratory experiments [7, 8]. Similarly, experiments with foraging animals have demonstrated risk-seeking behavior when expected reward in the environment is below average, further highlighting the role of contextual expectations in shaping subjective value [9].

A standard assumption is that a single reference point is updated gradually over time as new outcomes are observed. However, this cannot be the whole story, for several reasons. First, the decision making literature is scattered with observations that people can adopt multiple reference points [10, 11, 8, 12, 13], and recent work has shown that contextual cues can induce rapid alternation between multiple reference points [13]. Kahneman [14] likened the mental co-existence of reference points to ambiguous images (e.g., the Necker cube): the mind does not settle on one or average them together, but instead entertains them all in a state of tension. Second, evidence suggests that some patterns of irrational choice observed in behavioral experiments can be accounted for by biases in perception [15]. Given the pervasiveness of these choice anomalies both inside and outside of the lab, many have called for the extension of economic theory to explicitly allow for perceptual distortions arising from the way in which the brain encodes and retrieves information [16, 17]. Lastly, evidence from other domains suggests that humans organize their knowledge into discrete units (chunks, clusters, contexts, etc.) based on statistical regularities [18, 19, 20, 21, 22, 23], and thus it seems plausible that a similar form of “structure learning” might be invoked to organize the distribution of outcomes into discrete contexts.

In this paper, we pursue this idea theoretically and empirically. Following prior work [2, 3], we posit that reference points reflect rational expectations updated based on recent outcomes. However, we additionally assume that reference points can be discovered *de novo* via structure learning. Crucially, whereas prior research manipulates reference dependent choice through use of explicit contextual cues (e.g., [13]), here alternation between multiple reference points reflects inferences about the generative distributions to which different observations belong. Adapting a paradigm for studying structure learning in perceptual judgment [18], we present experimental evidence that risk preferences for gambles with outcomes drawn from a fixed distribution are influenced by the distribution of other gambles experienced in the same context (varied across blocks), which serve as implicit contextual cues. Crucially, if the fixed and variable distributions are sufficiently different, then the contextual effects are attenuated, indicating that they were assigned to distinct reference points. This attenuation effect is eliminated when the variable distribution is changed gradually across blocks, suggesting that a single reference point is applied to both distributions when their differences are made less salient. These patterns are captured by a Bayesian structure learning model of reference point formation.

## Results

### Experiment 1

Participants completed a binary choice task (see Materials and Methods for details) in which they made choices between a certain lottery (e.g., 50 points) and a lottery offering a fair chance of doubling or forfeiting the same amount (e.g., 50% chance of 100 points, 50% chance of 0 points). While prior experimental and theoretical work suggests that preferences are nonlinear in probabilities [14], we make the simplifying assumption that individuals’ preferences are linear in probabilities [2, 3]. Thus, the expected values of the two options were equivalent. Unbeknownst to participants, the expected value was randomly sampled from one of two Gaussian distributions (denoted A and B; Figure 1). Distribution A was held fixed across all conditions, whereas the mean of distribution B was varied. Each block featured an equal number of trials drawn from A and B, and the two trial types were visually indistinguishable.

**Figure 1:**
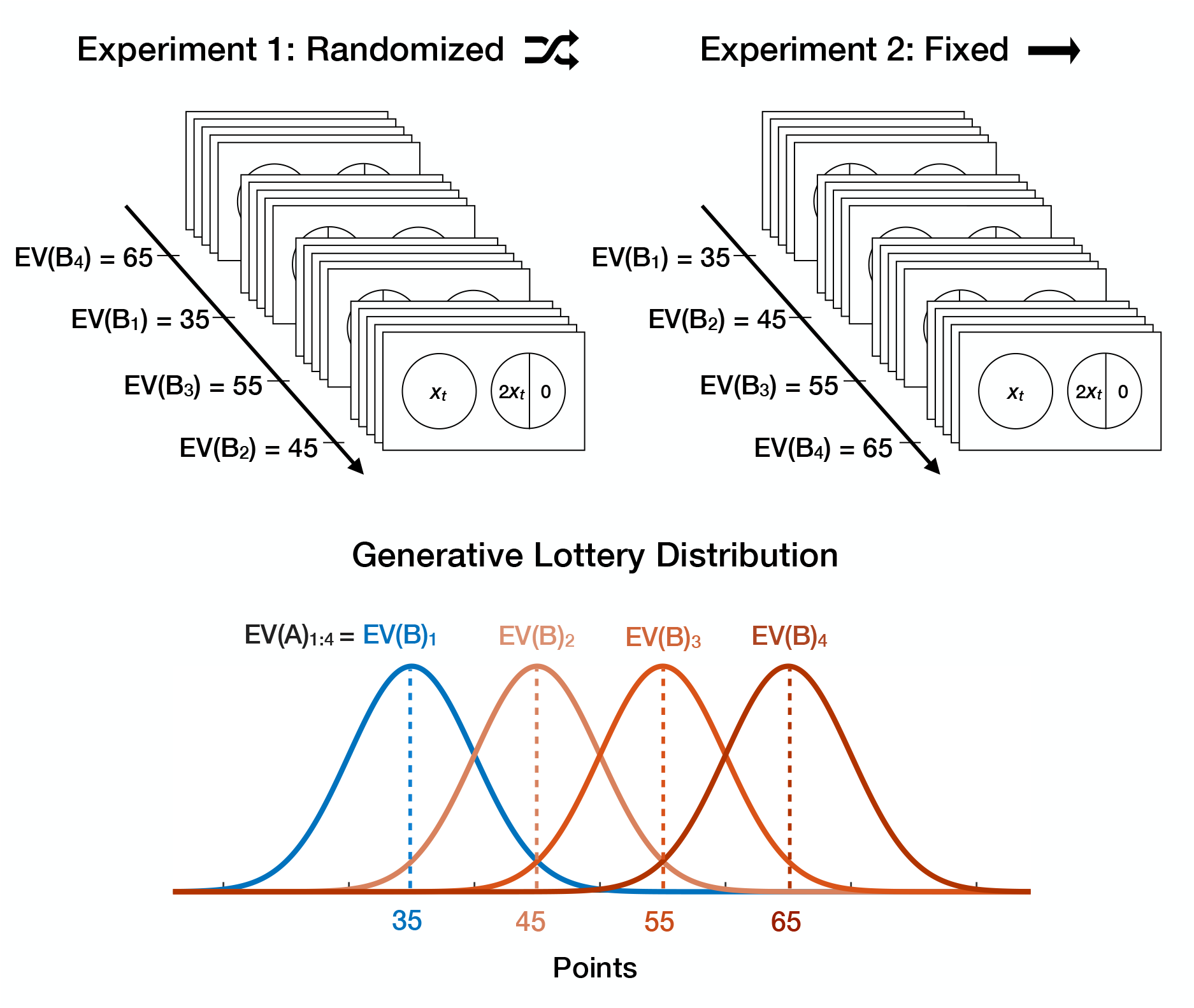
Task Design. On each trial, participants made a choice between a certain reward (displayed as a circle containing the reward amount in points) and a lottery with the same expected value (displayed as a divided circle containing twice the certain reward amount on one half and 0 on the other half). Expected values were drawn from one of two Gaussian distributions (A or B). Distribution A was fixed across all 4 conditions, whereas the mean of distribution B varied across conditions. In Experiment 1, the order of conditions was randomized across blocks; in Experiment 2, the conditions were ordered such that the mean of B increased monotonically across blocks.

According to our structure learning account, which we formalize below, participants should cluster A and B trials together when the means of the distributions (i.e., EV(A) and EV(B)) are close, because this is a statistically parsimonious account of the data. In this case (Figure 2A), a single reference point is applied to both A and B trials, corresponding to the expectation of the merged distribution. Since the model allows the latent distribution to drift slowly over time, sufficiently small changes in the mean of B impact the reference point applied to both A and B trials. More specifically, the reference point for A trials should increase with the mean of B, as long as the two trial types are clustered together (Figure 2B). Crucially, the model also predicts that participants should assign A and B trials into separate clusters when EV(A) and EV(B) are sufficiently distinct (Figure 2C). In this case, separate reference points are formed for A and B trials, corresponding to the expectations of each respective distribution.

**Figure 2:**
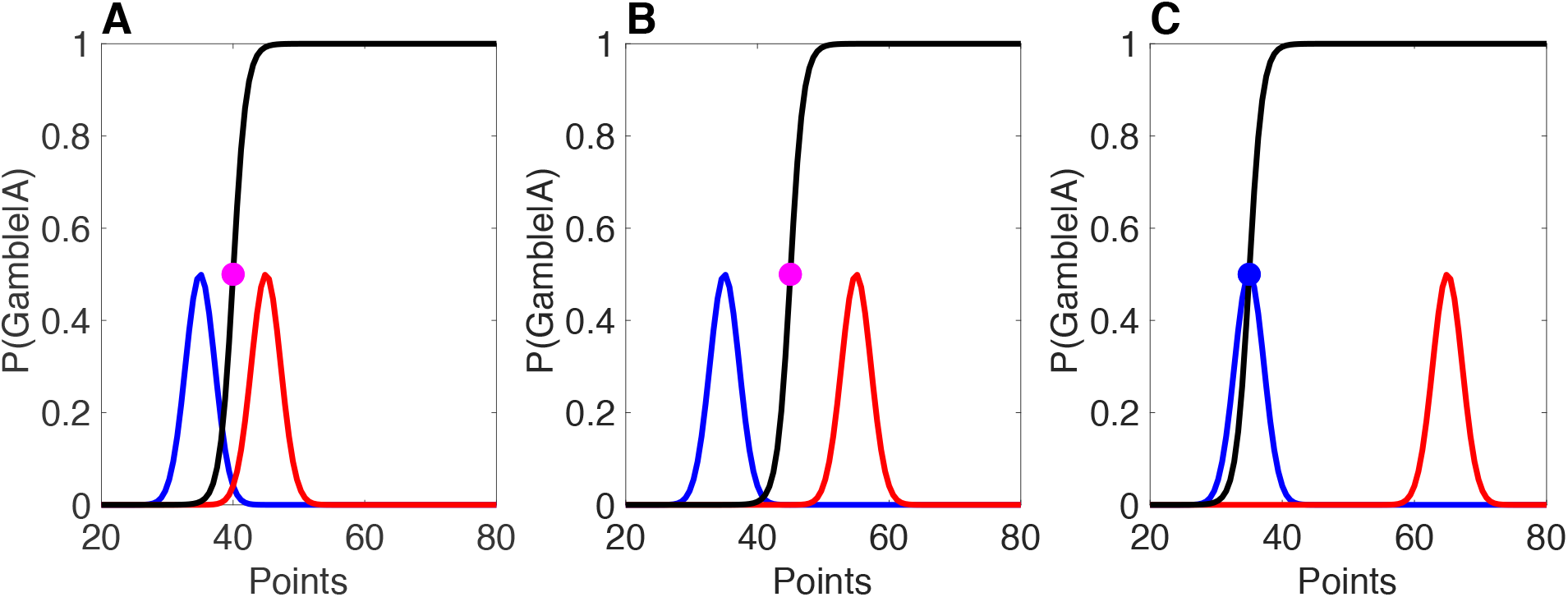
Reference point formation as structure learning. The black curve shows the probability of gambling on A trials, with risk-aversion above the reference point (indicated by a circle), and risk-seeking below the reference point; for risk-seeking participants, this pattern flips. The distribution of expected values for A trials is shown in blue, and the distribution for B trials is shown in red. (A) When the two distributions are close together, a shared reference point (purple circle) is formed based on the merged distribution. (B) For a moderate increase in the mean of the B distribution, the shared reference point increases. (C) For a sufficiently large increase in the mean of B, separate reference points are formed for A and B trials.

By design, the design of the task allows us to discern evidence for this shifting reference point by measuring how the probability of choosing the risky option (i.e., P(Gamble|A)) changes as a function of EV(B). Since EV(A) is fixed throughout the task, we can attribute any changes in P(Gamble|A) across conditions to changes in the reference point due to shifts in EV(B). The direction in which P(Gamble|A) is expected to change varies from person to person. For individuals who gamble more often as EV(A) increases relative to the reference point, P(Gamble|A) should mimic the U-shaped trajectory of EV(A) detailed above, but for those who prefer to gamble less as EV(A) increases relative to the reference point, P(Gamble|A) should follow an inverted U-shape.

In economics, this idiosyncrasy arises from differences in the curvature of the utility function, which maps objective outcomes (e.g., points or money) in terms of their subjective value (SV) [17]. This psychophysical perspective follows Bernoulli’s observation that risk aversion could be rationalized by diminishing marginal returns (i.e., concave utility). Thus, economists describe individuals who gamble less frequently with increasing value as “risk-averse,” and those who gamble more often as value increases as “risk-seeking” (see [24] for a review). A common misconception is that these labels capture people’s predispositions towards gambling, but it is not uncommon for a putatively “risk-averse individual to gamble more frequently than an individual characterized as “risk-seeking”. To minimize confusion, here we refer to the tendency to gamble more frequently with increasing SV as *positive value-dependent risk preference*, and we refer to the tendency to gamble less frequently as SV increases (or equivalently, to gamble more frequently as SV decreases) as *negative value-dependent risk preference*. To determine the appropriate set of predictions for each participant, we estimated the effect of expected value on the probability of gambling (see Methods for further detail). For individuals who gamble more often as EV(A) increases relative to the reference point, P(Gamble|A) should mimic the U-shaped trajectory of EV(A) detailed above, but for those who prefer to gamble less as EV(A) increases relative to the reference point, P(Gamble|A) should follow an inverted U-shape.

We tested these predictions in a data set of 92 participants (54 exhibited negative value-dependent risk preferences, 38 exhibited positive value-dependent risk preferences; see Materials and Methods for a description of how this designation was determined). The mean of B differed in increments of 10 across experimental conditions: 35, 45, 55, and 65 points, with the lowest value equal to the mean of distribution A. The order of these conditions was pseudo-randomized across blocks. We reasoned that randomization would make the difference between conditions highly salient. Note that, by chance, a small subset of participants were designated to complete the conditions in gradual order, but these participants were considered part of Experiment 2.

As shown in Figure 3, both participants with positive value-dependent risk-preferences and negative value-dependent risk-preferences changed their frequency of gambling for A trials (i.e., P(Gamble|A) non-monotonically as a function of EV(B). In accordance with our predictions, participants with negative value-dependent risk preferences first increased their risk preference and then decreased it, whereas participants with positive value-dependent risk preferences did the opposite. Note that, if the expectations-based reference point were based solely on an estimate of overall average reward, we would expect P(Gamble|A) to increase monotonically. Alternatively, if the reference point were based solely on the average reward for A trials, then we would expect no change across conditions. Thus, our experimental results provide strong support for our structure learning hypothesis.

**Figure 3:**
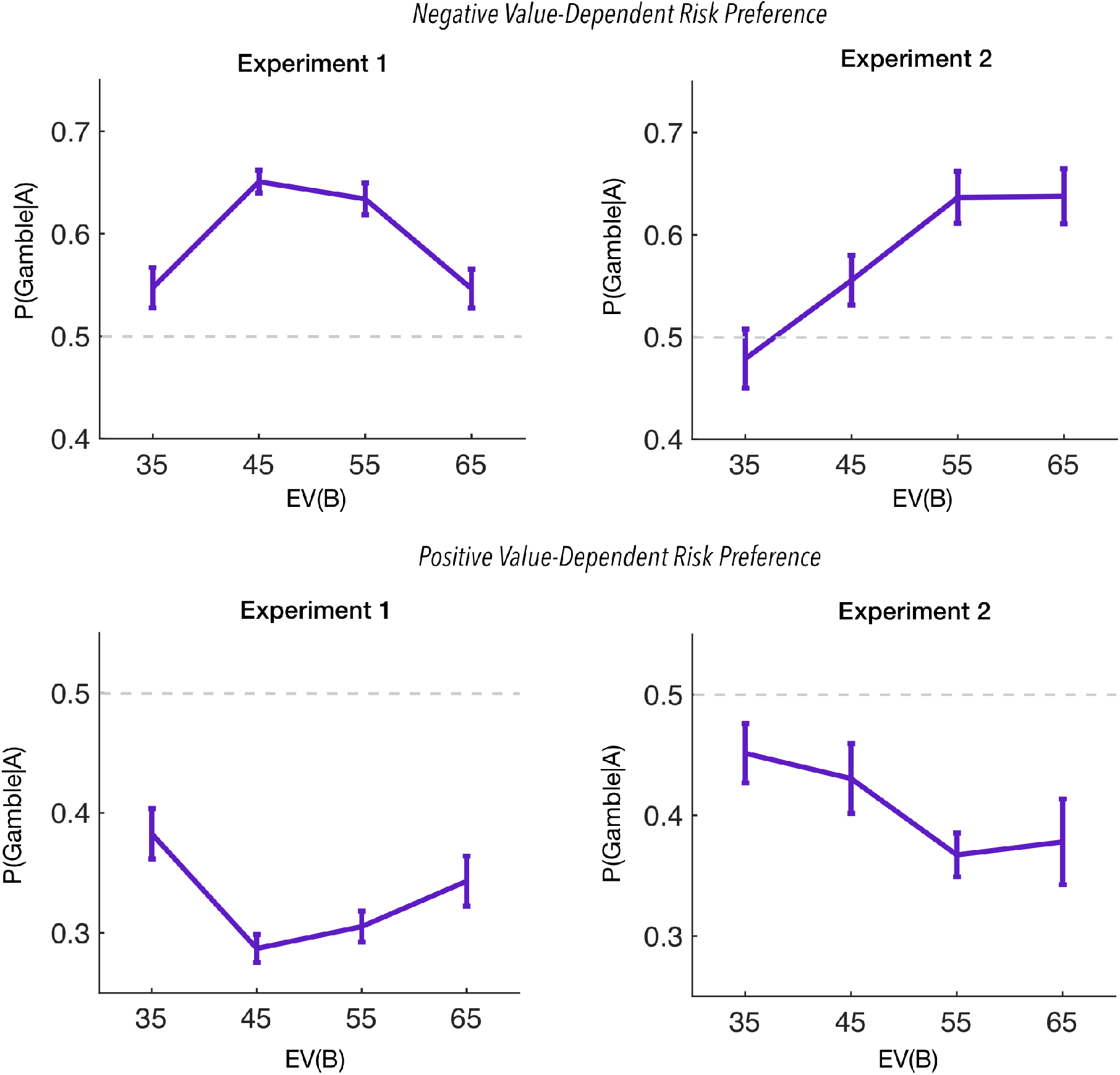
Probability of gambling (choosing the risky option) on lotteries drawn from distribution A, plotted as a function of the mean of distribution B. Results are shown separately for participants with negative value-dependent risk preferences (top) and participants with positive value-dependent risk preferences (bottom). Recall that risk preferences are not defined by the overall frequency of choosing the risky option, but rather, according to whether risk taking increases or decreases as a function of expected value (see Materials and Methods for details). Left: Experiment 1 results (mean of B randomized across blocks). Right: Experiment 2 results, where the mean of B increases monotonically across blocks. Error bars represent within-participant standard error of the mean.

To evaluate these patterns quantitatively, we fit a mixed-effects logistic regression model with the form P(Gamble|A) ~ Int. + EV(B) + EV(B)^2^, which allows us to capture quadratic effects of EV(B). We compared this to a model that lacked the quadratic term, and hence can only capture linear effects of EV(B). In both models, the identity of the participant was treated as a random effect, and parameters were estimated separately for the subset of participants with positive value-dependent risk preferences and for participants with negative value-dependent risk preferences.

The regression results are summarized in Table 1 (see Table S1 in the Supporting Information for the linear model results). For participants with negative value-dependent risk preference, the quadratic model fit yielded significant positive linear and negative quadratic coefficients. For participants with positive value-dependent risk preferences, the quadratic model fit yielded significant negative linear and positive quadratic coefficients. The quadratic effects constitute quantitative support for the non-monotonic pattern shown in Figure 3. For both participants with positive value-dependent risk preferences and participants with negative value-dependent risk preferences, likelihood ratios tests allowed us to reject the linear model relative to the quadratic model (negative value-dependent risk preference: Λ = 58.98, *p* < 4.8*e* − 12; positive value-dependent risk preference: Λ = 36.68, *p* < 2.1*e* − 07). Moreover, two standard model comparison metrics (Akaike information criterion and Bayesian information criterion) favored the quadratic model (Table 2).

**Table 1:** Parameter estimates for mixed-effects logistic regression model of P(Gamble|A), the probability of choosing the risky option when the expected value is drawn from distribution A. *β* = regression coefficient; SE = standard error; Int. = intercept; p-value = probability of observing data under the null hypothesis (based on t-statistic); EV(B) = expected value of distribution B.

| | | $\beta$ | SE | p-value |
| --- | --- | --- | --- | --- |
| **Experiment 1** | *Negative Value-Dependent Risk Preference* |  |  |  |
|  | Int. | -0.67 | (0.24) | 5.6e-03 *** |
|  | EV(B) | 1.1 | (0.20) | 1.4e-08*** |
|  | EV(B)<sup>2</sup> | -0.22 | (3.6e-02) | 1.1e-09*** |
|  | *Positive Value-Dependent Risk Preference* |  |  |  |
|  | Int. | 0.44 | (0.22) | 5.6e-02 * |
|  | EV(B) | -1.33 | (0.29) | 1.4e-06*** |
|  | EV(B)<sup>2</sup> | 0.25 | (5.5e-02) | 1.1e-06*** |
| **Experiment 2** | *Negative Value-Dependent Risk Preference* |  |  |  |
|  | Int. | -0.69 | (0.46) | 0.13 |
|  | EV(B) | 0.67 | (0.34) | 4.8e-02* |
|  | EV(B)<sup>2</sup> | -0.08 | (6.7e-02) | 0.23 |
|  | *Positive Value-Dependent Risk Preference* |  |  |  |
|  | Int. | 0.19 | (0.32) | 0.54 |
|  | EV(B) | -0.50 | (0.44) | 0.26 |
|  | EV(B)<sup>2</sup> | 0.066 | (9.4e-02) | 0.48 |

**Table 2:** Model comparison metrics for regression analyses. AIC = Akaike Information Criterion; BIC = Bayesian Information Criterion.

|  |  | **AIC** |  | **BIC** |  |
| --- | --- | --- | --- | --- | --- |
|  |  | Quadratic | Linear | Quadratic | Linear |
| **Experiment 1** | *Negative Value-Dependent Risk Preference* | 6703.7 | 6754.7 | 6763.0 | 6787.6 |
|  | *Positive Value-Dependent Risk Preference* | 4141.1 | 4169.7 | 4197.2 | 4200.9 |
| **Experiment 2** | *Negative Value-Dependent Risk Preference* | 2147.6 | 2144.2 | 2197.2 | 2171.7 |
|  | *Positive Value-Dependent Risk Preference* | 2438.7 | 2461.2 | 2489.7 | 2489.5 |

### Experiment 2

The results of Experiment 1 indicate that large, abrupt changes in distributional statistics can drive reference point formation. Based on findings from Pavlovian [25], motor [26], and perceptual [27, 28, 19] learning experiments, we hypothesized that subtler changes would obscure the differences between distributions and thus prevent new reference points from being formed. To test this hypothesis, we used the same conditions as in Experiment 1, but ordered them monotonically across blocks, such that each transition between blocks was associated with a small increase in the mean of distribution B. This gradual ordering features three 10pt shifts as opposed to the randomized ordering which guarantees at least one 20pt shift.

Data from 39 participants (18 with negative value-dependent risk preferences, 21 with positive value-dependent risk preferences) were analyzed using the same procedure described for Experiment 1. Consistent with our hypothesis that gradual change would obscure the distributional differences between conditions and lead to a single shared reference point, participants with negative value-dependent risk preferences increased their gambling on A trials monotonically as a function of EV(B), while participants with positive value-dependent risk preferences decreased their gambling monotonically (Figure 3).

Mixed-effects logistic regression confirmed this observation statistically. In contrast to the results of Experiment 1, we found no evidence for quadratic effects in either participant group (Table 1). Quantitative model comparison metrics favored the linear over the quadratic model (Table S2), and parameter estimates (summarized Table S1) showed significant linear effects of EV(B). Taken together, these results support our claim that a shared reference point tracked the gradually increasing EV(B) across blocks.

### Computational modeling

To account for experimental results, we generalize the notion of expectation-based reference point updating to incorporate structure learning. Following seminal work by Kőszegi and Rabin [2, 3], we assume that subjective utility has both reference-dependent and reference-independent components, and that reference points reflect contextual expectations based on a recency-weighted average of prior outcomes. However, whereas the model of Kőszegi and Rabin [2, 3] focuses on “how people react to departures from a posited reference point”, our primary endeavor is to provide a rigorous characterization of what the reference point is and where it comes from. Recent theoretical work has made foundational progress towards this goal by formalizing reference dependence as a product of Bayesian inference over features which serve as contextual cues. However, this work implicitly endows the learner with infinite representational capacity, despite evidence of limitations and information compression documented in psychology and neuroscience. By contrast, our work takes inspiration from the growing literature on rational inattention, which seeks to integrate these constraints with extant microeconomic theory using a rational approach [29].

Our main point of departure is the idea that the reference point corresponds to an agent’s expectation [2, 3, 4], rationally updating over time using Bayes’ rule and a nonparametric prior that allows the agent to cluster its experience into distinct latent causes [25, 19]. The expectation (and thus the reference point) can sometimes “jump” rather than adapt slowly. Most importantly for present purposes, structure learning can give rise to multiple reference points within the same context.

We develop the computational model in three stages. First, we describe the hypothetical data-generating process that characterizes the agent’s internal model of the world. Second, we formalize the inference problem facing the agent: to form beliefs about structure (latent causes) and the distribution of rewards associated with each latent cause. Equating reference points with these beliefs is the key innovation of this model. Third, we describe how beliefs are translated into a choice policy. We then fit the model to our data and show that it can capture the key phenomena observed in our experiments.

Let *x_t_* ∈ ℝ denote the reward payoff for the certain option (or equivalently the expected value of the risky option) on trial *t*. This reward is drawn from a Gaussian distribution with mean 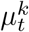 and variance *σ*^2^, where *k* = *z_t_* indicates the latent cause responsible for trial *t*. We assume a Gaussian prior over the initial condition 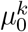, with mean 0 and standard deviation 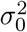. To allow for relatively small, gradual changes in the payoff distribution over time, we assume that the mean follows a Gaussian random walk: 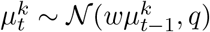, where *w* ∈ [0, 1] is a decay parameter that controls the rate of mean-reversion.

To allow for larger, abrupt changes in the payoff distribution, we assume that the latent cause can change over time, either by resampling an old latent cause or sampling a new latent cause. Thus, sufficiently small changes in the payoff distribution are perceived as gradual drifts in the mean of a single latent cause, whereas large, abrupt changes are more likely to be attributed to a new latent cause. Following previous work on structure learning [25, 19], we model the prior over latent causes with a Chinese restaurant process (CRP) [30, 31], which generates assignments of trials to latent causes according to the following sequential stochastic process:

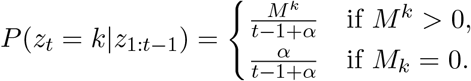

where *M ^k^* is the number of trials assigned to latent cause *k* up to trial *t*, and *α* ≥ 0 is a parameter controlling the number of latent causes. When *α* = 0, all trials are assigned to the same latent cause, and in the limit *α* → ∞, all trials are assigned to different latent causes. More generally, the expected number of latent causes after *t* trials is *α* ln *t*.

The computational problem facing the agent at time *t* is to infer the joint posterior over latent causes and their associated expected reward, as stipulated by Bayes’ rule:

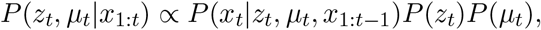

where the index 1: *t* indicates the set of all trials from 1 to *t*. The likelihood *P* (*x_t_|z_t_, μ_t_, x*_1:*t−*1_) and priors *P* (*z_t_*)*P* (*μ_t_*) are given by the generative process described above. Details about tractably approximating the posterior can be found in the Supporting Information.

We assume that preferences follow a reference-dependent quadratic utility function:

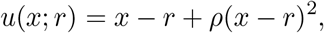

where *r* denotes the reference point (see below) and *ρ* controls the curvature of the utility function. In our experimental task, the expected utility of option *c* ∈ {certain, risky} is then given by:

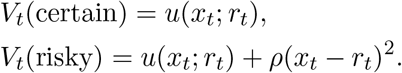

The curvature parameter *ρ* controls value-dependent risk preferences: *ρ* < 0 implies concavity for payoffs above the reference point (risk aversion or negative value-dependent risk preferences) and convexity for payoffs below the reference point (risk seeking or positive value-dependent risk preferences), as in Prospect Theory [1]. This pattern reverses for *ρ* > 0, and *ρ* = 0 implies neutrality. The quadratic utility function can also be understood as a special case of a mean-variance choice model [32, 33].

Under an expectations-based reference point model, *r* corresponds to the expected payoff 𝔼[*x*].

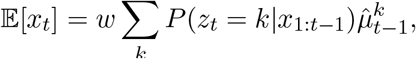

where 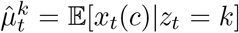 denotes the posterior mean payoff for latent cause *k*. In other words, the expectation on a given trial is the probability weighted average of the reward associated with each of its potential latent causes (see Supporting Information).

To allow for some stochasticity and bias in choice, we model the choice policy with a logistic sigmoid function *f* (*v*) = 1/(1 + *e^−v^*):

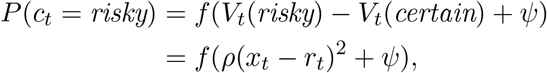

where *c_t_* is the choice on trial *t*, and *ψ* models an overall bias for gambling (*ψ* > 0) or choosing the certain option (*ψ* < 0).

In summary, the model assumes that observations arise from latent causes whose parameters change gradually over time. Agents reason backward from observations to latent causes using Bayes’ rule. Small changes in the distribution of observations are attributed to a single latent cause, whereas abrupt changes are attributed to switches between latent causes. Each latent cause is linked to choice behavior through an expectations-based reference point (the mean of the latent cause’s distribution over payoffs), such that agents have different risk preferences for payoffs that are greater or less than the mean.

The data from Experiments 1 and 2 were fit with two versions of the structure learning model (see Materials and Methods for model-fitting procedures): the full model (*α* > 0) that can learn multiple reference points, and a restricted model (*α* = 0) that learns a single reference point. Figure 4 shows the gambling probabilities for both models. The full model is able to capture the key findings: (1) a non-monotonic risk preference on A trials as a function of EV(B) in Experiment 1; (2) a monotonic risk preference in Experiment 2; and (3) opposite patterns of modulation for risk-averse and risk-seeking participants.

**Figure 4:**
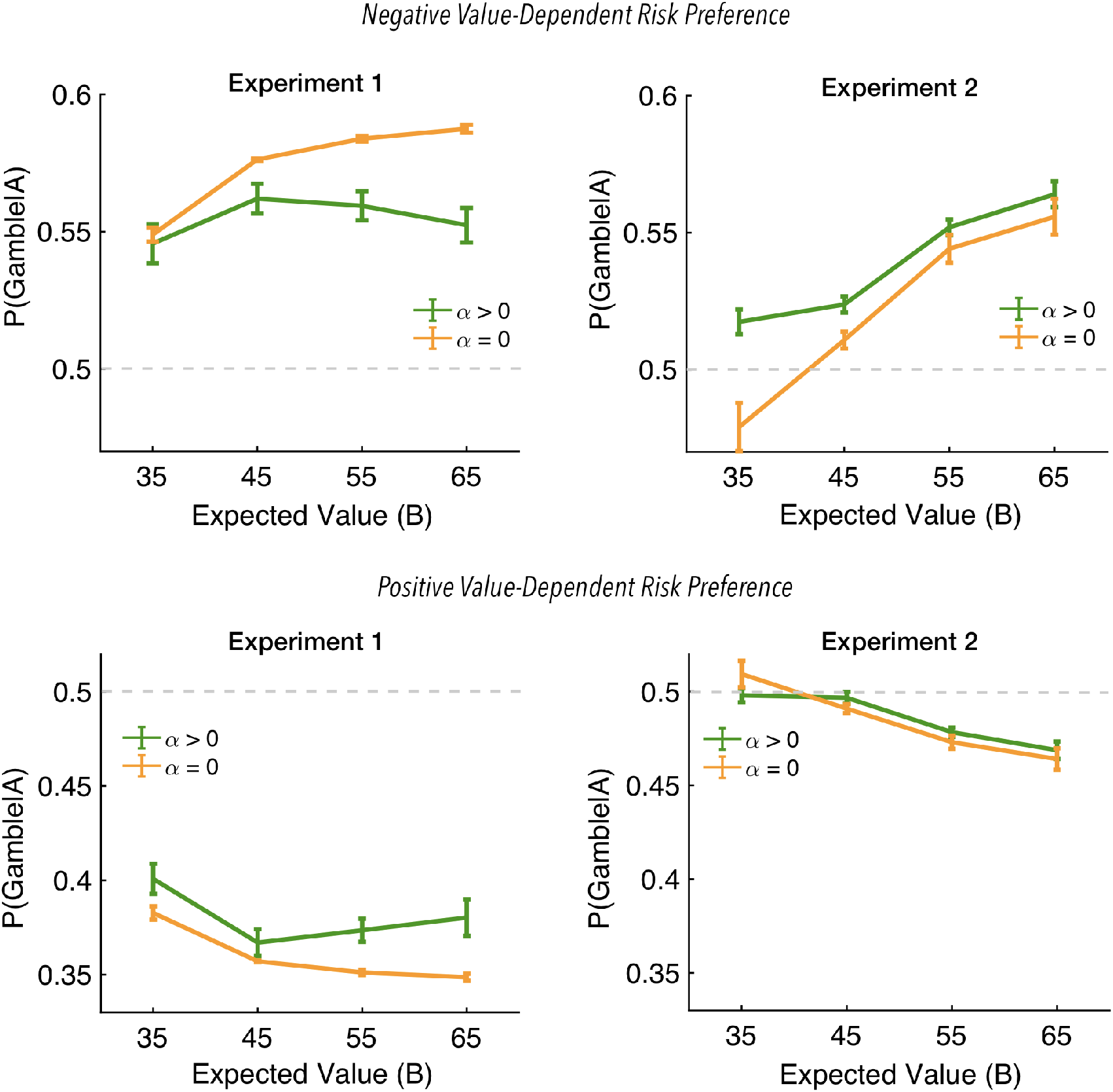
Model fits for the probability of gambling (choosing the risky option) on lotteries drawn from distribution A, plotted as a function of the mean of distribution B. Results are shown separately for participants with positive value-dependent risk preferences (top) and participants with negative value-dependent risk preferences (bottom). The *α* > 0 curve shows the fit of the full structure learning model that adaptively infers new reference points, and the *α* = 0 curve shows the fit of the restricted model in which all trials are forced to use the same reference point. Left: Experiment 1 results (mean of B randomized across blocks). Right: Experiment 2 results, where the mean of B increases monotonically across blocks. Error bars represent within-participant standard error of the mean.

The critical feature of the model is the ability to segregate A and B into separate latent causes when their expected values are perceived as sufficiently different (see supplemental methods for a trial-by-trial illustration). The importance of this feature is highlighted by the fact that the restricted model is unable to capture the non-monotonic risk preference in Experiment 1. The direct correspondence between these effects and the structural inferences of our clustering algorithm suggests that our results are robust to the particular parameterization of the utility function. Indeed, although we specify a quadratic value function for the practical purpose of analyzing the empirical data, in the supplement, we show that the same pattern of results can be achieved by using a Prospect Theory [1] utility function with reference points identified by structure learning.

## Discussion

Reference points play a fundamental role in theories of decision making, yet where they come from and how they change with experience has been an enduring puzzle. The research reported here sheds light on these questions, demonstrating that reference points arise from inferences about latent structure. The key idea is that new reference points are created when the prize distribution undergoes a dramatic change. In contrast, reference points are updated incrementally when the prize distribution undergoes gradual change.

The structure learning account of reference point formation makes a non-trivial prediction, which we confirmed experimentally: whereas small changes in the mean of the prize distribution can shift the reference point up or down, thereby altering risk preferences, large changes will actually have a *diminished* effect due to the creation of a new reference point. In other words, the magnitude of change affects risk preference non-monotonically. This pattern holds both for participants with positive value-dependent risk preferences as well as participants with negative value-dependent risk preferences. Importantly, when the mean changes gradually across blocks of the experiment, risk preference changes monotonically, in accordance with our prediction that gradual change will lead to reference point updating rather than the creation of a new reference point.

Our model of reference point formation is a significant generalization of the expectations-based reference point model [2, 3]. The core idea remains the same—reference points reflect expectations— but allows different expectations to form in different contexts. The notion that reference points are context-dependent has been widely acknowledged in psychology, but without a systematic formal treatment like the one proposed here [10, 34, 11, 8, 14].

A strong claim of our theory is that the brain tracks multiple reference points across time, invoking different reference points in a context-dependent manner. Brain imaging could be used to obtain independent evidence for this claim by identifying a neural correlate of the reference point, which could then be used to predict variability in risk preferences. Functional MRI studies have exploited this idea for a fixed structure [35, 36, 37], but have not yet investigated the role of structure learning.

While we have focused on expectations about prizes, the same logic can be applied to other economic variables, such as probabilities and delays. One interesting question is whether reference points apply to the joint space of economic variables, or whether these variables are dissociable, with reference points forming and updating independently. Either scenario could be formalized in our modeling framework, and could be addressed experimentally by orthogonally manipulating the magnitude of changes in different variables simultaneously.

Another important direction for future research is extending the framework to multi-attribute decisions, where much of the experimental and theoretical research on reference points has focused (e.g., [38, 39, 40, 41]). In principle this extension is straightforward, by modeling each latent cause with a multivariate distribution over attributes [19]. Further modifications might utilize additional parameters of the latent context [42] or even allow for latent contexts to relate hierarchically such that multiple reference points might be applied to a single decision simultaneously [34].

In summary, our findings implicate an important and hitherto unappreciated role for structure learning in decision making. These findings dovetail with results in a diverse set of domains, including memory [19, 43], social cognition [20], categorization [44], and perception [45], all of which involve some form of structure learning. Indeed, our experimental design closely mirrored previous experiments on perceptual judgment [18]. A common modeling framework, based on nonparametric Bayesian inference, can explain many aspects of behavior across these different domains, suggesting that there may be a set of basic computational principles that govern structure learning in the brain.

## Materials and Methods

### Participants

We recruited 200 individuals to complete the experiment online using the Amazon Mechanical Turk (MTurk) service. Participants had to correctly answer questions to ensure comprehension of the instructions before proceeding. In addition to $2 base pay, each participant was awarded a bonus payment based on the realization of a randomly selected trial. Points were converted into dollars such that the minimum bonus was $1 and the maximum was $2. Participants gave informed consent, and all procedures were approved by Harvard Universitys Institutional Review Board.

### Exclusion criteria

In line with recommendations for studies conducted using Amazons Mechanical Turk (AMT) service, careful instructions and a priori exclusion criteria were applied to ensure data quality (Crump et al., 2013). Of the 186 subjects who progressed to the end of the experiment, fourteen were excluded for failing to provide responses for greater than 20% of trials overall or greater than 20% of trials within a given block (i.e., more than 40 trials overall trials or more than 10 trials within a block). Given that participants had ample time to respond to each question (up to 30 seconds), we reasoned that these subjects were not devoting sufficient attention to the task. By contrast, we found that the remaining subjects completed a minimum of 92.5% of trials and all but 5 achieved a completion rate of 98% or higher. Thus, although the cut-off rate of 80% completion (20% omission) was determined *a priori*, any rate ≤ 92.5% completion would have resulted in the same final set of subjects. Another 27 subjects were excluded for choosing the left vs. right or risky vs. certain lottery on over 90% of trials. These biases correspond with preferences that fall outside of the range we can reasonably capture using this paradigm. Gambling on less than 10% (or more than 90%) of trials leaves at least five of the eight (2 distributions x 4 conditions) sub-conditions with only 2/25 trials on which the subject deviated from the preferred option. Figure S1 shows that the tails of the distribution of gambling frequencies for the final sample follows a normal distribution whose tails correspond with these boundaries. Finally, 17 subjects were excluded because they showed no effect of expected value on choice behavior. Due to the fixed structure of the gamble relative to the certain gain, the only aspect of the task that varied across trials was expected value. Thus, if subjects are engaged in the task, risk taking should vary as a function of expected value (see Identification of value-based risk preferences for further details). Our overall exclusion rate of 29% is well in line with the findings of research on quality control measures on MTurk [46][47][48].

### Procedure

Participants completed 200 trials in which they chose between a certain option (guaranteed x points) and a risky option (2x points with probability 0.5). The side of the screen on which the risky option appeared was counterbalanced across trials. Participants were given a relatively unlimited amount of time (up to 30 seconds) to make each choice (subject to the 15 minute deadline for completion of the task). Participants did not receive feedback about the outcome of their choice. Once a response was recorded, the task transitioned to a 1.5 second inter-trial interval during which a fixation cross appeared. To discourage the use of simple heuristics (e.g., always choosing the risky lottery gamble) and promote sustained attention, catch trials were embedded randomly within each block. On these trials (4 total across the entire experiment), the options were mismatched so that either the certain or risky option had twice the expected value of the other.

### Design

Experiment 1 consisted of 4 conditions (50 trials per condition) presented in random order across participants. Each condition differed only in the mean of B, which took on values of 35, 45, 55, or 65 points. Distribution A had a fixed mean of 35 points across all conditions. The variance of both distributions was equal to 5 points. Experiment 2 was almost identical to experiment 1, except that the conditions were presented in order of increasing mean.

### Identification of value-dependent risk preferences

In order to distinguish participants with positive value-dependent risk preferences from those with negative value-dependent risk preferences, we estimated the effect of expected value on the probability of gambling separately for each participant. For this analysis, we focused exclusively on the condition in which EV(A) = EV(B) = 35. Here A and B trials should be clustered together with a reference point corresponding to their estimated mean. In this case, subjective value is a roughly a linear function of objective, expected value. By contrast, when multiple reference points are applied, SV is a non-monotonic function of EV. For example, consider a condition with reference points *z* and 2*z* where *z* > 1. If value is purely reference dependent, then an outcome with an expected value of *z* + 1 will have a positive subjective value, whereas an outcome with an expected value of 2z-1 would have no subjective value. As A trials and B trials were indistinguishable in this condition, we used all 50 trials from this condition More specifically, we fit a logistic regression model to participants choice data as a function of expected value *x_t_* on trial t, *P* (*c_t_* = *risky*) = *f* (*x_t_*), where *f* () is the logistic sigmoid function with intercept *β*_0_ and slope *β*_1_. Participants with negative regression coefficients were assumed to have negative value-dependent risk preferences, and participants whose regression coefficient was significantly positive were considered to have positive value-dependent risk preferences. As mentioned in the exclusion criteria, participants whose regression coefficient was insignificant were not included in further analyses.

### Model fitting

Each participant’s choice data were fit separately using maximum likelihood estimation of parameters. For the full structure learning model, the free parameters were *α*, *ψ*, *ρ*, *Q*, and *σ*^2^. For the restricted model, the free parameters were *ψ*, *ρ*, *Q*, and *σ*^2^ (with *α* fixed to 0). The remaining parameters were fixed as follows for both models: 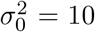 and *w* = 1.0. Numerical optimization was used to find maximum likelihood estimates of the free parameters, with 5 random initializations to avoid local optima.

### Model comparison

We used two standard metrics for model comparison: the Akaike Information Criterion (AIC) and the Bayesian Information Criterion (BIC).

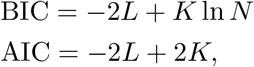

where *L* is the maximum likelihood value, *K* is the number of parameters, and *N* is the number of data points. Both metrics balance model fit (*L*) against model complexity (*K*), but the BIC penalizes complexity more strongly.

## Acknowledgements

S.J.G. was supported by the National Institutes of Health (CRCNS R01-1207833) and the Office of Naval Research (N000141712984).

## Supporting Information (SI)

### Posterior approximation

As described in the main text, the agent observes the expected payoff *x_t_* ∈ ℝ on trial *t* and uses this observation to update her belief about the latent cause *z_t_* responsible for the observation, as well as her belief about the mean of the payoff distribution 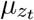 associated with the latent cause. Here we provide the details of how the update is computed.

The posterior over *z_t_* is given by:

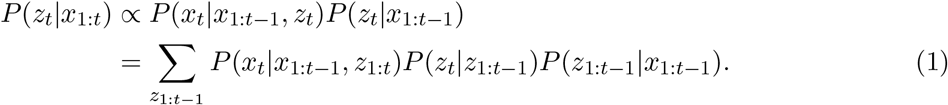

Because the sum over *z*_1:*t−*1_ is computationally intractable, we use a “local” maximum *a posteriori* (MAP) approximation [44, 49, 19], which replaces the marginalization with a maximization:

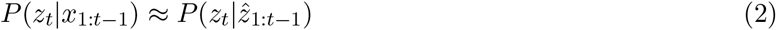

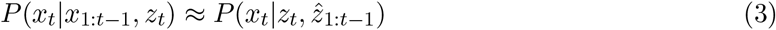

where *P* (*z_t_|ẑ*_1:*t−*1_) is the Chinese restaurant process (CRP) and *ẑ*_1:*t−*1_ is defined recursively according to:

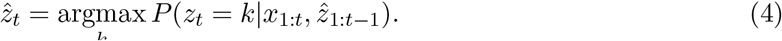

The local MAP approximation does not in general yield the history of latent causes with the highest posterior probability, because it does not update past assignments after observing new information. However, it is often sufficiently accurate, as attested by its use in machine learning applications [50].

Using the local MAP approximation, the likelihood is given by:

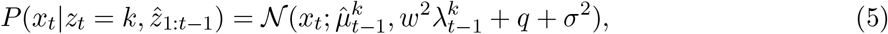

where *w* ∈ [0, 1] is a decay parameter, *q* is the diffusion noise variance, *σ*^2^ is the observation noise variance, 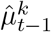 is the posterior mean for cause *k* after observing trials 1 to *t* − 1, and 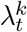 is the posterior variance. The mean and variance are updating according to the Kalman filtering equations:

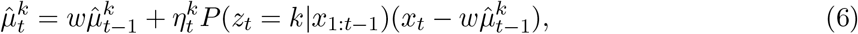

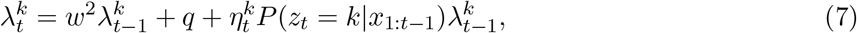

where 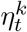 is the Kalman gain (learning rate), given by:

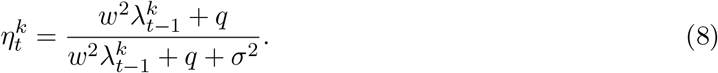

### Prospect Theory as an alternative value function

The reference points identified by the structure learning model can serve as input to a variety of utility functions beyond the standard quadratic value function presented in the main text. Given the expected value of an outcome *x* and the reference point *r*, Prospect Theory defines subjective value or utility of perceived gains (*x* ≥ *r*) and perceived losses (*x* < *r*) according to a piece-wise power function. Our simplified variant:

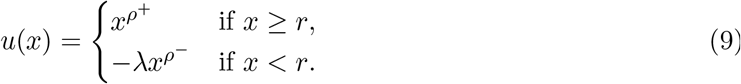

where loss aversion coefficient *λ* represents the multiplicative weighting of perceived losses relative to perceived gains, and the exponents *ρ*^+^ and *ρ*^−^ control the degree of curvature of the utility function for perceived gains and losses respectively. When *ρ* < 1 the utility function exhibits diminishing sensitivity to changes in value as the absolute value increases and thus is concave for relative gains and convex for relative losses; when *ρ* < 1, the utility function exhibits increasing sensitivity to changes in value as the absolute value increases, and thus is concave in the gain domain and convex for relative losses. For simplification, we assume the degree of curvature is symmetric for perceived gains and losses such that *ρ*^+^ = *ρ*^−^ = *ρ*. This constraint has been shown to provide a superior fit in prior empirical applications [51, 52].

**Figure S1:**
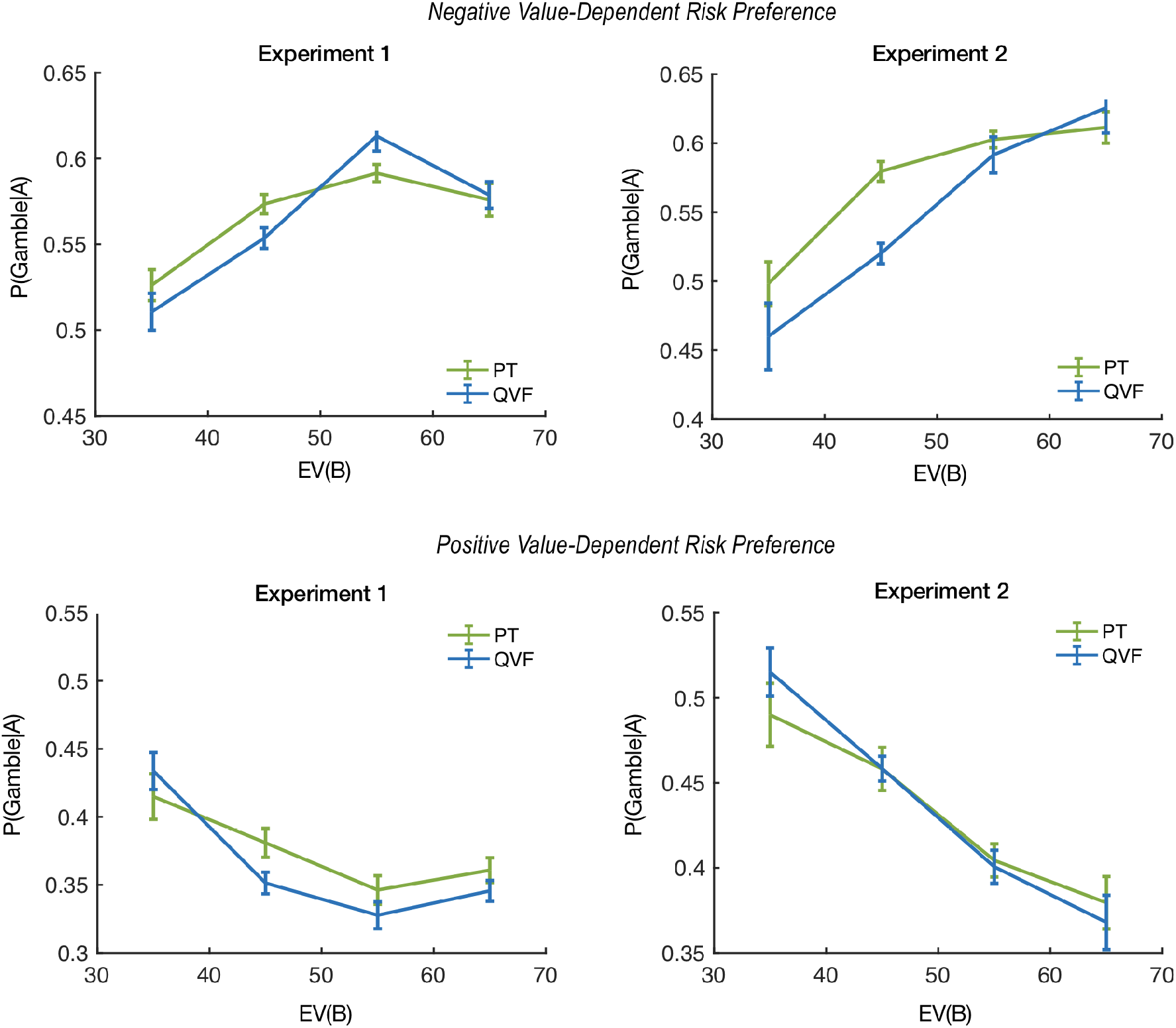
Model fits for the generic quadratic value function (QVF) and Prospect Theory (PT) the probability of gambling (choosing the risky option) on lotteries drawn from distribution A, plotted as a function of the mean of distribution B. Results are shown separately for participants with negative value-dependent risk preferences (top) and participants with positive value-dependent risk preferences (bottom). The green curve shows the fit of the structure learning model with a Prospect Theory Value function, and the blue curves show the fit using a simple Quadratic Value Function (QVF). Left: Experiment 1 results (mean of B randomized across blocks). Right: Experiment 2 results, where the mean of B increases monotonically across blocks. Error bars represent within-participant standard error of the mean.

**Figure S2:**
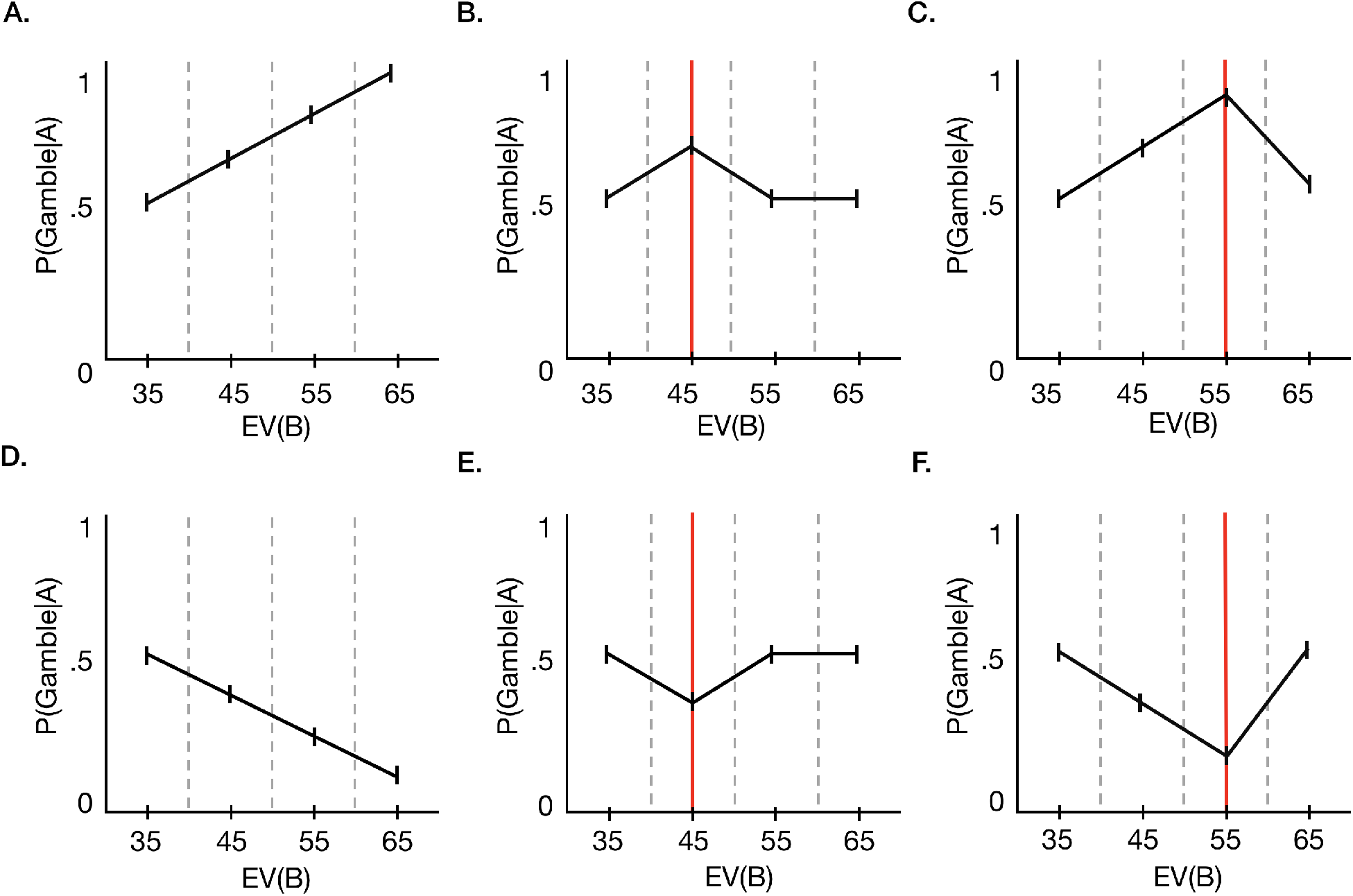
Idealized patterns of gambling frequency as a function of EV(B) for hypothetical groups of participants with different perceptions about where the perceived split between A and B occurs. Each column provides separate predictions for participants with positive value dependent risk preferences (A-C; upper panel) and participants with positive value dependent risk preferences (D-F; lower panel). Grey dashed lines demarcate the set of trials belonging to each condition and thereby indicate where shifts in the generative distribution of EV(B) occur. Solid grey lines, by contrast, indicate where the perceived split between A and B trials is assumed to occur.

**Table S1:**
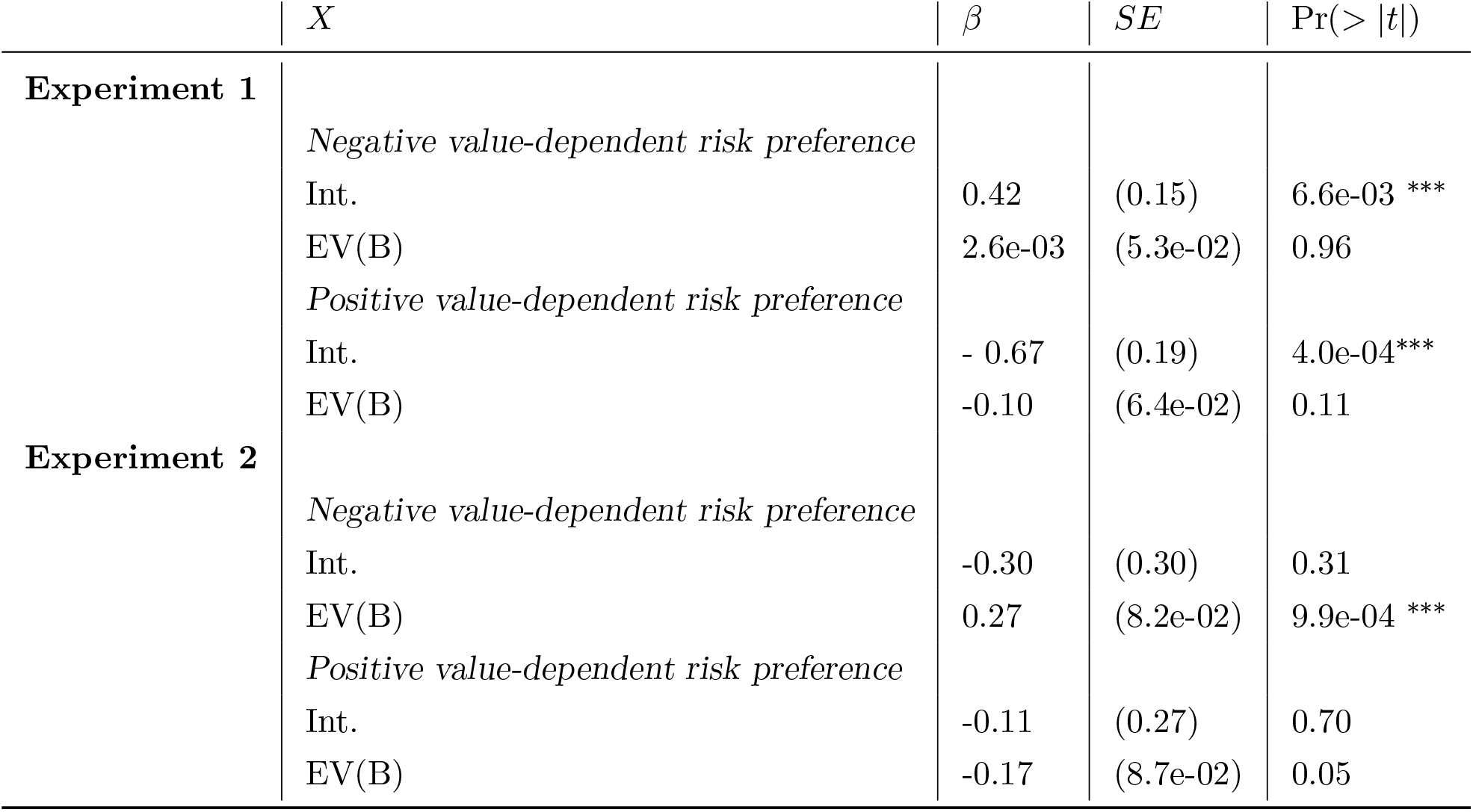
P(Gamble|A) as a linear function of EV(B). *β* = regression coefficient; SE = standard error; Int. = intercept; EV(B) = expected value of distribution B.

The data from Experiments 1 and 2 were fit with two versions of the structure learning model which differed only in the particular function used to map observations and reference points to utility: one used the quadratic utility function described in the full text while the other used the simplified version of Prospect Theory discussed above. Each participant’s choice data were fit separately using maximum likelihood estimation of parameters. For the structure learning component of both models, we assumed 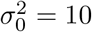 and *w* = 1.0, leaving three free parameters: *α*, *σ*^2^, and *q*. For the quadratic utility function, the two free parameters were the value-dependent risk preference *ρ* and the gambling bias *ψ*. As described above (Equation 9), the free parameters of the Prospect Theory utility function were risk-preference *ρ* and loss aversion *λ*. Numerical optimization was used to find maximum likelihood estimates of the free parameters, with 5 random initializations to avoid local optima. Figure S1 shows the gambling probabilities for both models. Both are able to capture the key findings: (1) a non-monotonic risk preference on A trials as a function of EV(B) in Experiment 1; (2) a monotonic risk preference in Experiment 2; and (3) opposite patterns of modulation for participants with negative value-dependent risk preferences and participants with positive value-dependent risk preferences.

### Model-free signatures of structure learning

The key prediction of our model is that participants in Experiment 1 will be more likely to form separate reference points for A and B trials than participants in Experiment 2. As described in the main text, the reference point for A trials increases with EV(B) when A and B trials are clustered together and then decreases when A and B trials are clustered separately. Therefore, in general, we expect P(Gamble|A) to change roughly as a linear function of EV(B) for a participant in Experiment 2 (Figure S2, panels A and D) and we expect P(Gamble|A) to change as a fairly (inverted) U-shaped function of EV(B) for a participant in Experiment 1 (Figure S2, panels B, C, E, and F). While these predictions are relatively straightforward at the participant level, formal assessment of these hypotheses at the group level is complicated by the fact that individuals differ not only by *whether* they perceive a split between A and B, but also by *where* they perceive a split between A and B.

In particular, idiosyncrasy in structure learning has two major consequences for our group-level results. First, although we would expect the group-level effects to be nonmonotonic in the extreme case in which all participants inferred a split during the same condition, here we simply expect the change in P(Gamble|A) as a function of EV(B) to be more (inverse) U-shaped, for participants in Experiment 1 than for participants in Experiment 2. Critically, by comparing Experiment 1 to Experiment 2, we control for other potential sources of quadraticity (e.g., due to diminishing marginal effects of shifts in EV(B). Secondly, heterogeneity in where the perceived split between A and B occurs is not only a random effect, but one that is expressly non-linear with respect to the fixed effect of EV(B). For this reason, we cannot simplify our predictions using a linear model or even a joint test of multiple linear models.

To better understand these points, consider cases where there is consensus regarding the condition in which A and B split. Logic dictates that the perceived split must occur within one of the three conditions where EV(B) is greater than EV(A), but we can disregard the particular trial on which the perceived split occurs without loss of generality. In each case, the hypothesized (inverse) U-shaped effect can be re-framed as a joint hypothesis about two simple linear effects. If the perceived split takes place when EV(B) = 45 (Figure S2, panels B and E), the hypothesized (inverse) U-shaped effect can be re-framed as a joint hypothesis about two simple linear effects: The first effect would be a significant change in P(Gamble|A) as EV(B) increased from 35 to 45, and the second effect would be a linear effect of EV(B) on P(Gamble|A) in the opposite direction across the three conditions where EV(B) = 45, 55, and 65 respectively. However, if the perceived split takes place when EV(B) = 55 or 65 (Figure S2, panels C and F), we expect an overall linear effect of EV(B) on P(Gamble—A) across the three conditions where EV(B) = 35, 45, and 55 and a significant difference in the opposite direction from 55 to 65. Since these expectations contradict one another, this approach has no efficacy in cases like ours in which the parsing of A and B trials is subjective.

By contrast, the quadratic, logistic mixed-effects regression model explicitly accounts for heterogeneity of this sort by treating individual differences in the quadratic coefficient (i.e., magnitude and direction) as a random effect. This offers a parsimonious approach to assessing our model-free hypotheses - a quadratic effect is not a sufficient condition of nonmonotonicity, but it is a necessary one. Accordingly, by demonstrating that the fixed-effects quadratic term is significant at the group level for Experiment 1 but not for Experiment 2, we corroborate the predictions of the model.

### Risk preferences vs. gambling biases

In the main text we emphasize the theoretical distinction between value-dependent risk preferences and gambling biases. In our structure learning model these variables are formalized by the parameters *ρ* and *ψ* respectively. Although each has a computationally unique influence on choice behavior, it is conceivable that these phenomena might be shaped, in part, by common cognitive mechanisms. However, there is as of yet relatively little research on whether (and if so, how) these variables relate to one another. Although our study was not designed to address this question explicitly, the interplay between these two aspects of risky choice in relation to the dynamic structure of our task and the divergent patterns of reference dependence in the two experiments highlights valuable issues for future research to address.

Consistent with a recent study by Rigoli et al., 2018 ([13]), we found no correlation between value dependent risk preferences and gambling propensity as measured by *ρ* and *ψ* (p*>*.1) overall. However, when each dataset is considered separately, this result only holds for Experiment 2 (p¿.1). In Experiment 1, there is a significant negative relationship between these parameters (r = -.32, p*<*.01). As an alternative to comparing the model parameters, we can also examine overall gambling frequency as a function of putative risk preference (see Identification of value-dependent risk preferences in Materials and Methods). For the pooled dataset, this analysis corroborates the afore-mentioned lack of correspondence between risk preference and gambling frequency approximated via *ρ* and *ψ*. Specifically, an unpaired two-sample t-test of the hypothesis that the two independent samples (i.e., participants with positive-vs. negative-value-dependent risk-preferences), came from distributions of gambling frequencies with equal means gambling frequency, indicated that the null hypothesis (”means are equal”) cannot be rejected at the 5 % significance level (*t*_129_ = 0.96, *p* = .33). Intriguingly, the fidelity of these results to those of the parametric approach did not hold at the experiment level. Instead, the results for both individual experiments mirrored the null finding for the pooled results (Experiment 1: *t*_90_ = −0.57, *p* = .57; Experiment 2: *t*_37_ = 1.55, *p* = .13).

At a glance, this appears at odds with the patterns of behavior shown in Figure 3 where P(Gamble|A) appears largely above .5 for participants with negative value-dependent risk preferences and largely below .5 for participants with positive value-dependent risk preferences. It is important to note here that these figures only depict A trials. Intuitively, it makes sense that participants with positive value-dependent risk preferences gamble less frequently on A trials in conditions where EV(A), is less than EV(B).

By the same logic, gambling frequency should be statistically indistinguishable on A and B trials when EV(A) = EV(B). Therefore, if gambling propensity is independent from value-dependent risk preference, we would expect P(Gamble—A, EV(B) = 35) not to differ as a function of risk preference. This is true for Experiment 2 where P(Gamble|A,EV(B)= 35) *≈ .*47 for both types of value-dependent risk preferences (Figure 3, right panel). However, for Experiment 1, P(Gamble—A) remains lower for participants with positive value-dependent risk preferences than for participants with negative value-dependent risk preferences even in the condition where EV(B) = EV(A). The key to making sense of this difference lies in the fact that participants in Experiment 2 always start the task with the condition in which EV(B) = 35, whereas the majority of participants in Experiment 1 first encounter a condition in which EV(B) is greater than 35. As a result of this difference, participants in Experiment 1 are more likely to have higher reference points leading into the condition where EV(B) = 35 than participants in Experiment 2. This explains why P(Gamble|A, EV(B)=35) is higher for participants with negative (vs. positive) value-dependent risk preferences but it also raises a new question. Since EV(A) = EV(B) in this condition, higher reference points also predict overall differences in risk taking (i.e., on both A and B trials)-how can this be given that gambling propensity is independent from value-dependent risk preference?

**Figure S3:**
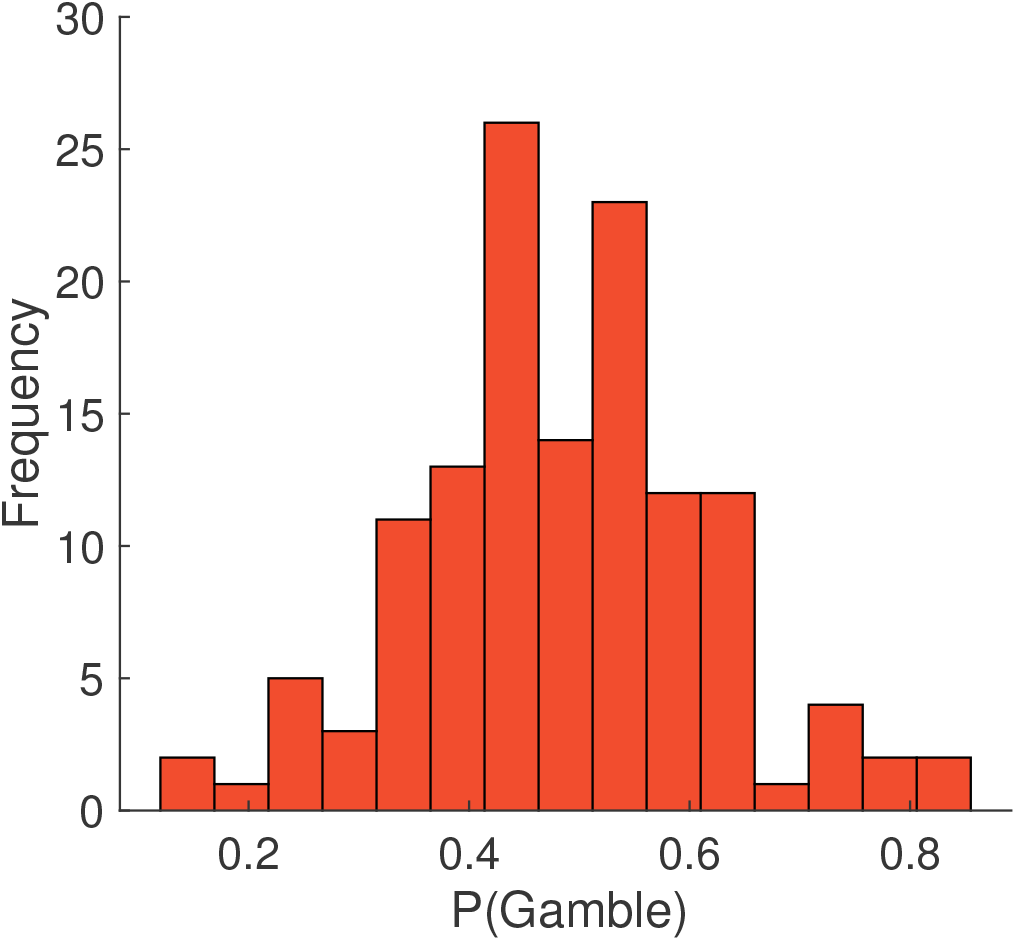
Histogram plotting the distribution of gambling frequencies (% of trials on which the risky option was chosen) for the final subject sample (*N* = 131).

The missing piece of the puzzle is illustrated in Figure S3. Although neither risk preference nor the Expected Value of B (EV(B)) predicts overall gambling frequency in Experiment 1 (*βEV* (*B*) = −4.5 *×* 10^−4^, *p* = .93), participants with negative value-dependent risk preferences are significantly more likely to gamble than participants with positive value-dependent risk preferences in the condition where EV(B) = 35 (*t*_90_ = 2.62, *p* = .01). In the left panel of Figure S3 we can see that in Experiment 1, there is a significant interaction between risk preference and EV(B) (Generalized linear mixed-effects model: ’*P* (*Gamble*) ~ 1 + *EV* (*B*)\**ρ̂*+ (1 + *EV* (*B*)\**ρ̂|participant*)’ where *ρ*̂ indicates putative value-dependent risk preference ; *β_EV_* _(*B*):*ρ̂*_ = .03, *p < .*001).

**Figure S4:**
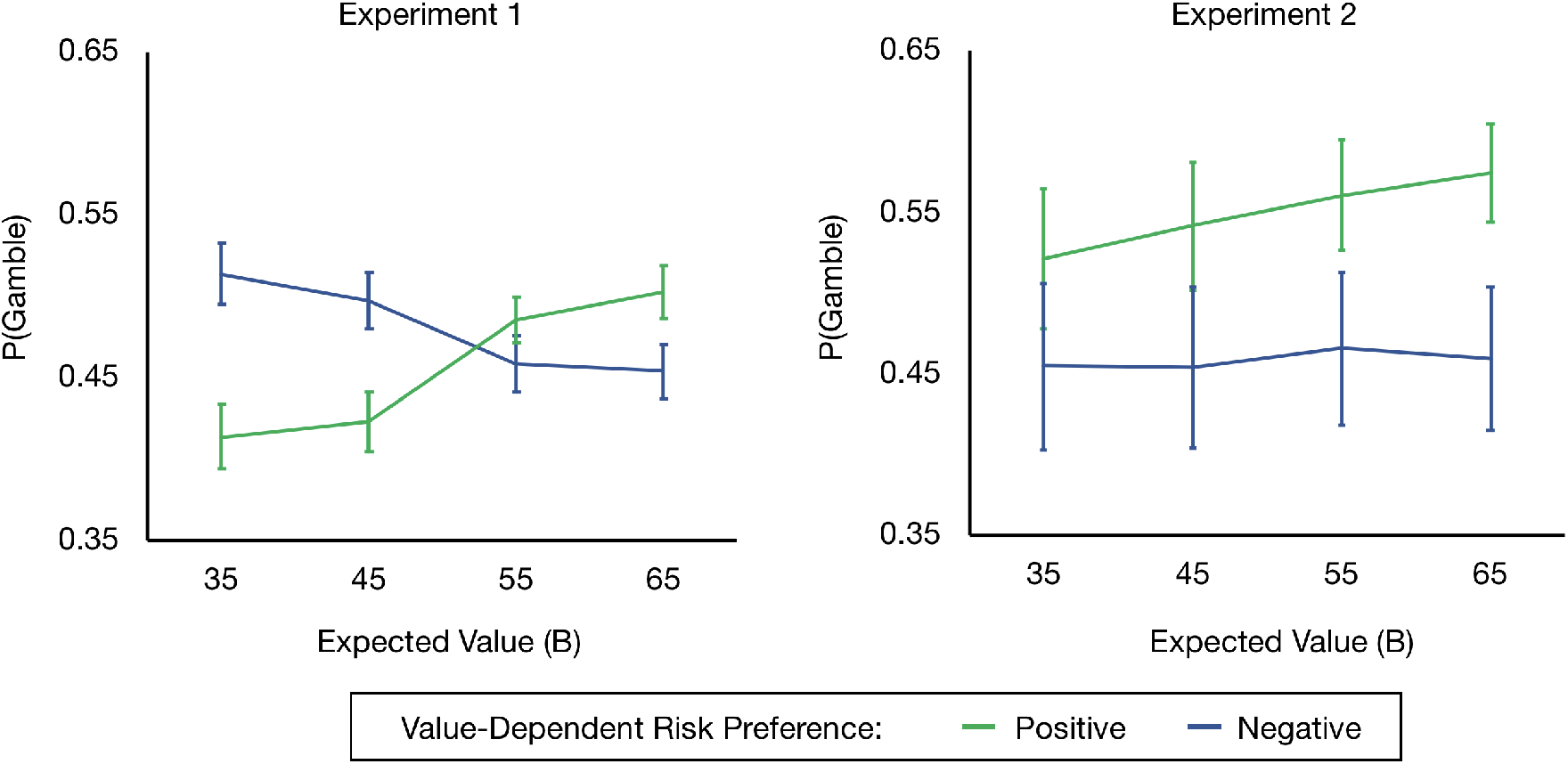
Average gambling frequency (both A and B trials) as a function of the Expected Value of B and value-dependent risk preference (Positive-in green; Negative-in blue) for Experiment 1 (left) and Experiment 2 (right). Error bars reflect the standard error of the mean. Between the two experiments, there is no significant difference in overall gambling frequency (*t*_129_ = −1.28, *p* = .20), and within each experiment, there is no significant difference in overall gambling frequency as a function of risk preference (Experiment 1: *t*_90_ = −0.57, *p* = .57; Experiment 2: *t*_37_ = −1.55, *p* = .13). Furthermore, overall gambling frequency does not change as a function of EV(B) (Generalized linear mixed-effects model: *P* (*Gamble*) ~ 1 + *EV* (*B*) + (1 + *EV* (*B*)*|participant*); Experiment 1: *βEV* (*B*) = −4.5 *×* 10^−4^, *p* = .93; Experiment 2: *βEV* (*B*) = 1.1 *×* 10^−3^, *p* = .42). However, within Experiment 1, there is evidence of a significant interaction between risk preference and EV(B) that is not present in Experiment 2 (Generalized linear mixed-effects model: *P* (*Gamble*) ~ 1 + *EV* (*B*)\**ρ̂*+ (1 + *EV* (*B*)\**ρ̂|participant*) where *ρ̂* indicates putative value-dependent risk preference ; Experiment 1: *β_EV_* _(*B*):*ρ̂*_ = .03, *p* < .001^***^; Experiment 2: *β_EV_* _(*B*):*ρ̂*_ = 7.6 *×* 10^−4^, *p* = .13).

**Figure S5:**
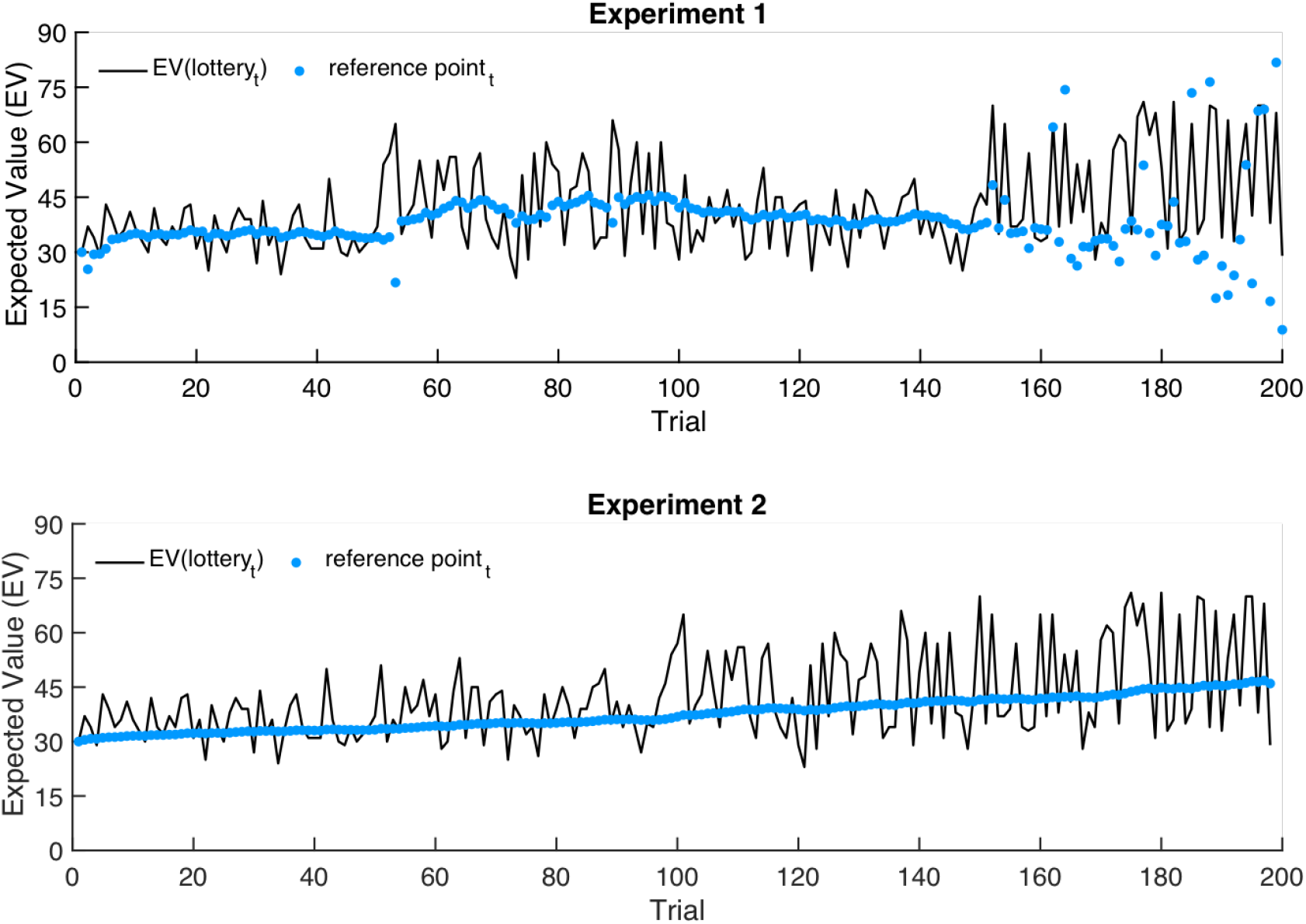
Schematic of trial-by-trial model predictions simulated for Experiment 1 (upper panel) and Experiment 2 (lower panel). Observed values on each trial are shown in black while the model estimates (i.e., reference points) are shown in blue.

**Figure S6:**
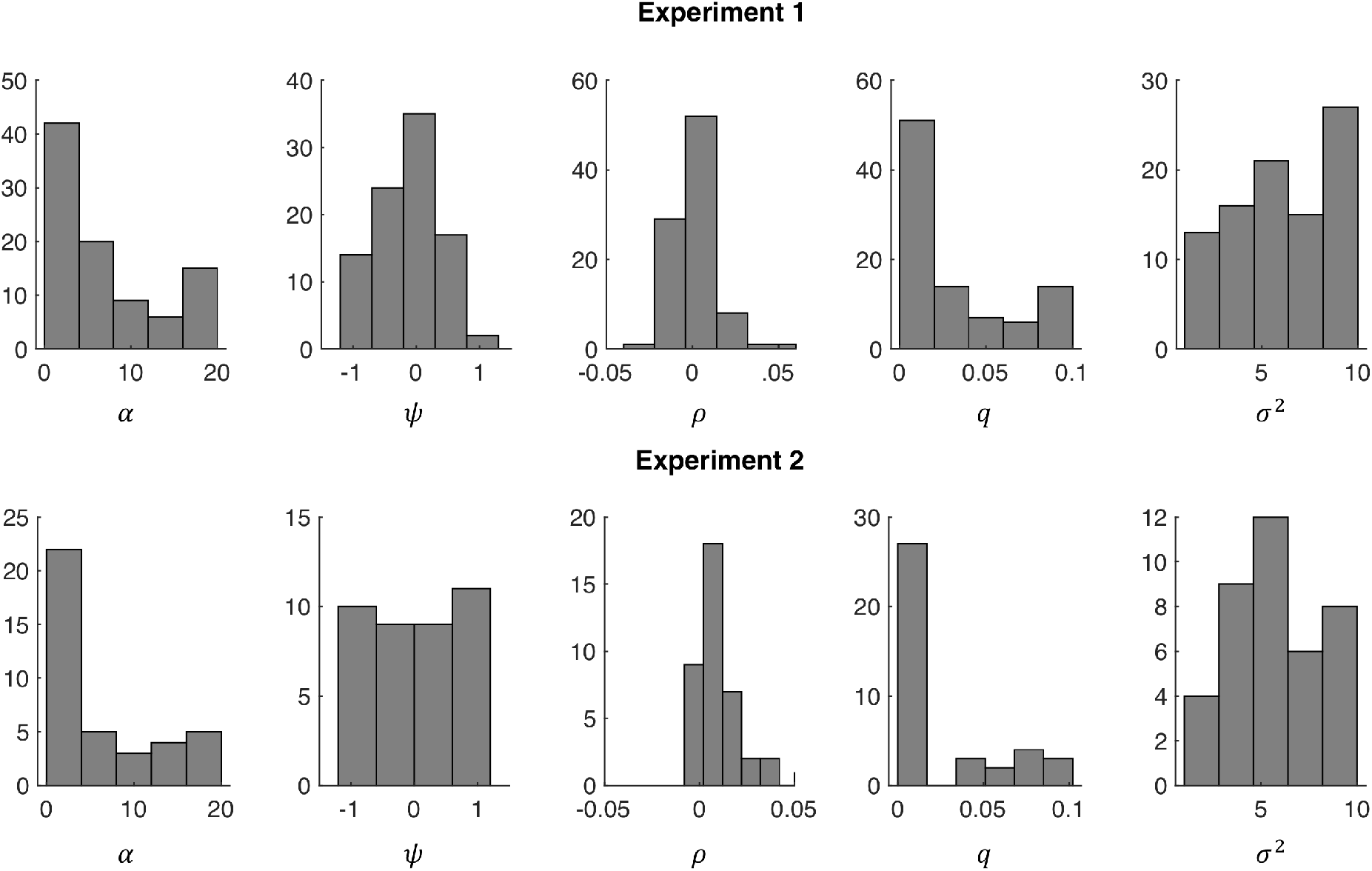
Distributions of individual fit values for the free parameters of the structure learning model for Experiment 1 (above) and Experiment 2 (below) (*N*_1_ = 92, *N*_2_ = 39, *N_total_* = 131).

